# The impact of comorbidity of intellectual disability on estimates of autism prevalence among children enrolled in US special education

**DOI:** 10.1101/011528

**Authors:** Andrew Polyak, Richard M. Kubina, Santhosh Girirajan

## Abstract

**Objectives:** While recent studies suggest a converging role for genetic factors towards risk for nosologically distinct disorders including autism, intellectual disability (ID), and epilepsy, current estimates of autism prevalence fail to take into account the impact of comorbidity of these neurodevelopmental disorders on autism diagnosis. We aimed to assess the effect of potential comorbidity of ID on the diagnosis and prevalence of autism by analyzing 11 years of special education enrollment data.

**Design:** Population study of autism using the United States special education enrollment data from years 2000-2010.

**Setting:** US special education.

**Participants:** We analyzed 11 years (2000 to 2010) of longitudinal data on approximately 6.2 million children per year from special education enrollment.

**Results:** We found a 331% increase in the prevalence of autism from 2000 to 2010 within special education, potentially due to a diagnostic recategorization from frequently comorbid features like ID. In fact, the decrease in ID prevalence equaled an average of 64.2% of the increase of autism prevalence for children aged 3-18 years. The proportion of ID cases potentially undergoing recategorization to autism was higher (p=0.007) among older children (75%) than younger children (48%). Some US states showed significant negative correlations between the prevalence of autism compared to that of ID while others did not, suggesting differences in state-specific health policy to be a major factor in categorizing autism.

**Conclusions:** Our results suggest that current ascertainment practices are based on a single facet of autism-specific clinical features and do not consider associated comorbidities that may confound diagnosis. Longitudinal studies with detailed phenotyping and deep molecular genetic analyses are necessary to completely understand the cause of this complex disorder. Future studies of autism prevalence should also take these factors into account.

**STRENGTHS AND LIMITATIONS:**

- We present a large-scale population study of autism prevalence using longitudinal data from 2000-2010 on approximately 6.2 million children enrolled in US special education.
- We provide one possible but compelling explanation for increase in autism prevalence and show that current ascertainment of autism is based on single facet of clinical features without considering other comorbid features such as intellectual disability
- We are not able to dissect the exact frequency of comorbid features over time as US special education allows for enrollment under only one diagnostic category and does not document comorbidity information.
- This study examines how comorbidity of related phenotypes such as ID can impact estimates of autism prevalence and does not assess other factors reported to impact autism prevalence.

## INTRODUCTION

Autism is a neurodevelopmental disorder characterized by impairments in social reciprocity, speech and communication, and restricted, repetitive and stereotyped patterns of behavior^1^. Several epidemiological reports have suggested an apparent increase in the prevalence of autism^2–4^. A recent study by the United States Center for Disease Control estimated the prevalence of autism among 8-year-old children, within the Autism and Developmental Disabilities Monitoring (ADDM) network sites in 2010, to be 1 in 68 children^5^. This estimate was a documented 123% increase in prevalence when compared with the data from 2002 (1 in 150 children). Another study, based on a population screening of 7-to 12-year-old elementary school children in a South Korean community in 2006, estimated an overall autism prevalence of 1 in 38 children^6^.

While the rise in autism prevalence has been attributed to various factors including increased awareness^7^ and broadening of the diagnostic criteria^3 4^, significant clinical heterogeneity and the non-specific molecular etiology of autism have precluded robust estimates of prevalence^8 9^. Further, there is a documented clinical overlap or comorbidity of nosologically distinct neurodevelopmental disorders^10 11^. For example, premorbid social impairment or pervasive developmental disorders (PDD) have been observed in 50-87% of individuals with childhood-onset schizophrenia^12–16^. Similarly, features of intellectual disability (ID) have been reported in as high as 68% of individuals with autism^17^, and epilepsy and attention-deficit hyperactivity disorders (ADHD) have been reported to co-occur in as high as 38.3% and 59%, respectively, of children with autism^9 18 19^. Accumulating evidence from genomic studies suggests a converging role for genetic factors towards a common molecular etiology for these varied neurodevelopmental disorders^12 20–22^. Although the epidemiological studies, to date, report statistically significant increases in the prevalence of autism, they fail to take into account the effect of the comorbidity of other disorders on the diagnosis and prevalence of autism. To better understand the effect of comorbid features on autism prevalence, we systematically analyzed 11 years (2000-2010) of epidemiological data on an average of 6.2 million children per year from the United States special education enrollment. We examined the frequency and age-specific prevalence of autism and frequently comorbid clinical features. Our results suggest that comorbidity of ID can significantly impact diagnosis and confound prevalence estimates of autism.

## METHODS

### Special education data

The Individuals with Disabilities Education Act (IDEA) is a law originally enacted in 1975 that ensures services to children with disabilities, ascertained by 13 disability categories throughout the United States. We obtained special education enrollment data, i.e., the number of children receiving services under various disability categories, from publicly available databases (IDEA part B) for the 50 US states. The IDEA part B database includes annual state-by-state counts of children aged 3-21 years documented by their age and classification of disorder (Figure S1). For the current study, we obtained enrollment data on approximately 6.2 million children per year aged 3-21 years over an 11-year period spanning 2000-2010 evaluated by special education and placed under one of the 13 disability categories (Table S1). Children recruited under the special education act are ascertained under only one disability category. However, reclassification to a different IDEA category is possible (e.g., an individual originally identified as having ID can be reevaluated and then reclassified into the autism category). Notably, school districts do not always use the *Diagnostic and Statistical Manual of Mental Disorders* criteria for classifying any of these diagnostic categories^23–25^. Further, the Department of Education’s legal definitions of disorders under the IDEA categories are generalized allowing for a broader interpretation, and therefore, the ascertainment for each disorder may vary between different states^4 26^.

### Data analysis

We used the United States intercensal estimates for ages 3-21 years for 2000 to 2010 to create proportions (number enrolled out of 10,000 children) for each ascertainment category. To assess the impact of comorbid features on autism prevalence, we estimated combined proportions of one or more related diagnostic categories including autism spectrum disorders (ASD), intellectual disability (ID), other health impairment (OHI), emotional disturbance (ED), and specific learning disability (SLD), whose phenotypes have significant comorbidity with autism^2 8 9 17^. We note that “developmental delay” was included within the intellectual disability category. Throughout the manuscript, the term “autism” is used interchangeably with “ASD”. We used simple linear regression to model the relationship between the year and prevalence of the phenotypic category. Using a nominal cut-off of p<0.05, we determined if the slope of each model was statistically significant.

## RESULTS

Consistent with recent reports^2 17^ we find a significant rise in the prevalence of autism from 1.2 per 1,000 in 2000 to 5.2 per 1,000 in 2010 (331% increase; linear regression, p=4.58×10^−10^) within the US special education population (Figure 1). However, significant decrease in prevalence from 8.3 per 1,000 in 2000 to 5.7 per 1,000 in 2010 was observed for ID (31% decrease, p=3.1×10^−10^). Significant decreases were also seen for categories of emotional disturbance (22%, p=6.1×10^−5^), and specific learning disability (19%, p=4.1×10^−8^) (Table S2). Based on recent studies suggesting common molecular etiology^10^ and documented reports of comorbidity of autism and other neurodevelopmental disorders^8^, we hypothesized that estimates of autism prevalence are confounded by the presence of comorbid phenotypes. We tested the impact of comorbidities on the prevalence of autism by first combining all related neurodevelopmental phenotypes whose prevalence changed significantly over the 11-year period. The combined prevalence of autism, ID, specific learning disability, and other health impairment did not change significantly from 2000 to 2010 (51.0 per 1,000 in 2000 to 50.3 per 1,000 in 2010; p=0.19) suggesting that the prevalence of autism is influenced by related comorbid phenotypes (Figure S2).

**Figure 1.**
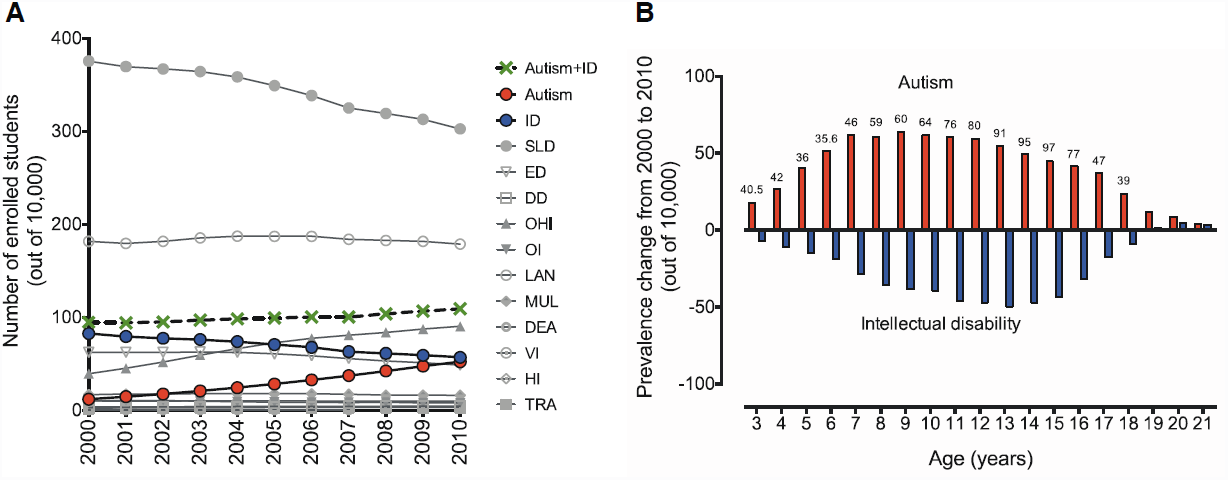
Prevalence of phenotypes from years 2000 to 2010 ascertained through special education enrollment. **(A)** Yearly prevalence (out of 10,000) is shown for the 13 special education categories including autism, intellectual disability (ID), specific learning disability (SLD), developmental delay (DD), other health impairments (OHI), emotional disturbance (ED), speech and language impairments (SLD), multiple disorders (MI), traumatic brain injuries (TRA), deaf-blindness (DB), deafness (DEA), orthopedic impairments (OI), hearing impairments (HI), and visual impairments (VI). The combined prevalence of autism and ID (Autism+ID) is also shown. US intercensal estimates for ages between 3 and 21 years were used as denominator in the prevalence calculations. **(B)** The age-specific changes (2000 to 2010) in prevalence of autism (red) and ID (blue) are shown from ages 3 to 21 years. Percentage of autism prevalence that can be attributed to diagnostic change from ID to autism is shown for each age as numbers above the red bars.

Interestingly, when only the prevalence estimates of autism and ID were considered together, the combined prevalence increased by 15% (9.5 per 1,000 in 2000 to 10.9 per 1,000 in 2010, p=1.7×10^−6^) (Figure 1A). This increase is 22-fold less than the increase seen for autism prevalence alone (331% vs. 15%), suggesting that a potential diagnostic recategorization from ID to autism can account for a significant amount of autism prevalence. In order to determine the ages at which these changes were most significant, we analyzed the age-specific changes in prevalence for autism and ID for the ages between 3 and 21 years. We found that an increase in autism prevalence, specifically across ages 3 to 18 years, corresponded to a decrease in the prevalence of ID (Figure 1B). These changes in autism prevalence compared to those in ID allowed us to calculate the possible magnitude of diagnostic recategorization from ID to autism. For example, at age 8 years, up to 59% of the increase in autism prevalence could be attributed to a diagnostic recategorization of ID. On an average, between the ages of 3 and 18 years the decrease in ID prevalence equaled 64.2% of the increase of autism prevalence. These estimates rise to as high as 97%, at age 15 years, with potential recategorization contributing to a 35-fold change in autism prevalence from 2000 to 2010 (Table S3). Further, older children (ages from 10 to 18 years) with ID were more likely to have a shift of diagnosis towards autism than younger children (ages from 3 to 9 years) (Mann Whitney test, p=0.007) (Figure S3). We also found that autism and ID can be distinguished based on age-prevalence. When evaluated by age, the prevalence of autism peaked between ages 7 and 9 years, while the age-specific prevalence for ID peaked between ages 11 and 19 years (Figure S4).

To assess if the autism diagnostic criteria are uniform across all US states, we compared the prevalence of autism with the prevalence of ID over the 11 years. We observed negative correlations, at significant levels, between the prevalence of autism compared to that of ID (Pearson’s correlation, r=−0.26, p=1.10×10^−9^) when all US states were considered together (Figure 2). Interestingly, these correlation estimates were higher for some US states than others (Table 1). US states with a higher prevalence rate for ID were more likely (Mann Whitney test, p=0.0476) to show a negative correlation with autism prevalence than those states with a lower prevalence of ID (Figure S5).

**Figure 2:**
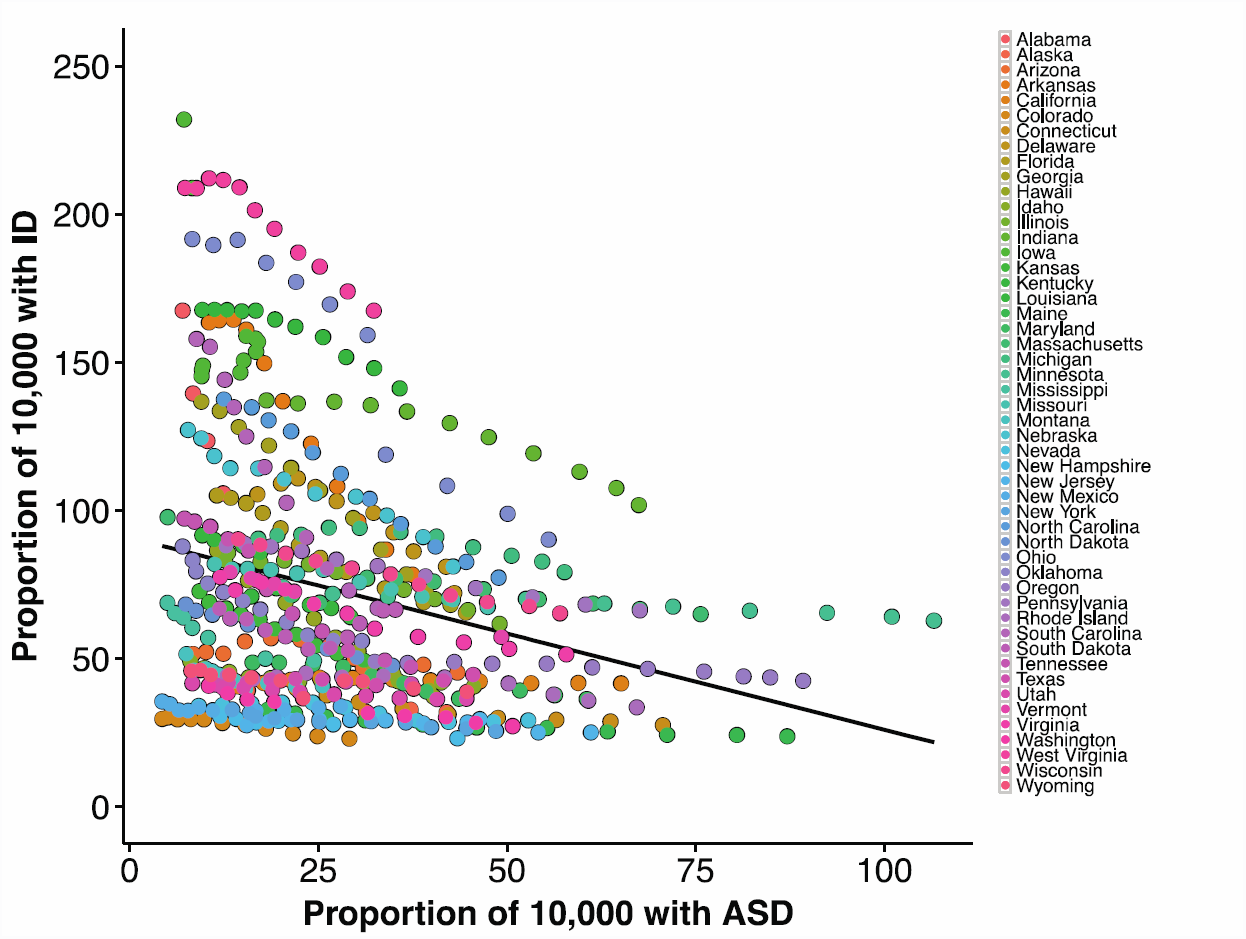
Negative correlation between the prevalence of autism to that of intellectual disability (r=-0.26, p=1.10×10^−9^). Pearson correlation coefficients were used to assess the relationship between the prevalence of autism and each of the comorbid phenotypic categories within the special education enrollment.

**Table 1:**
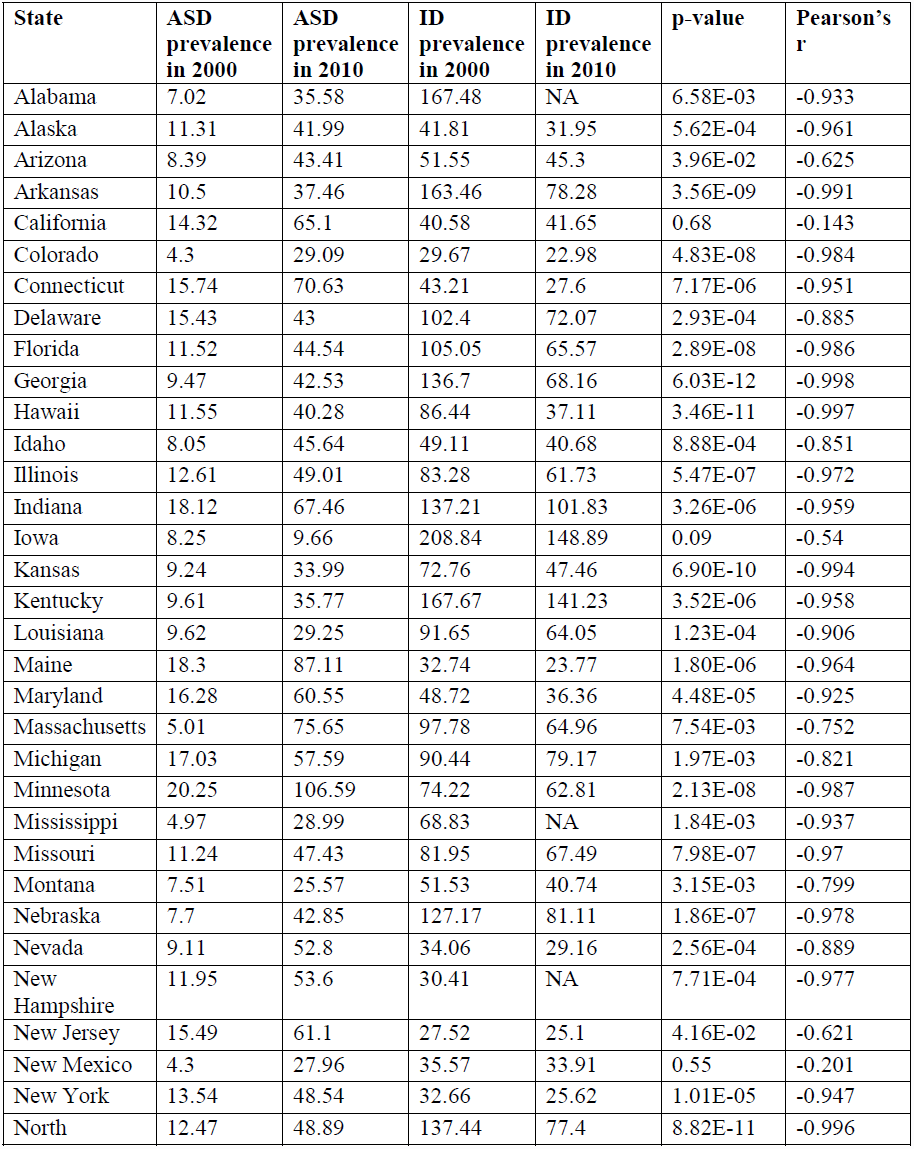
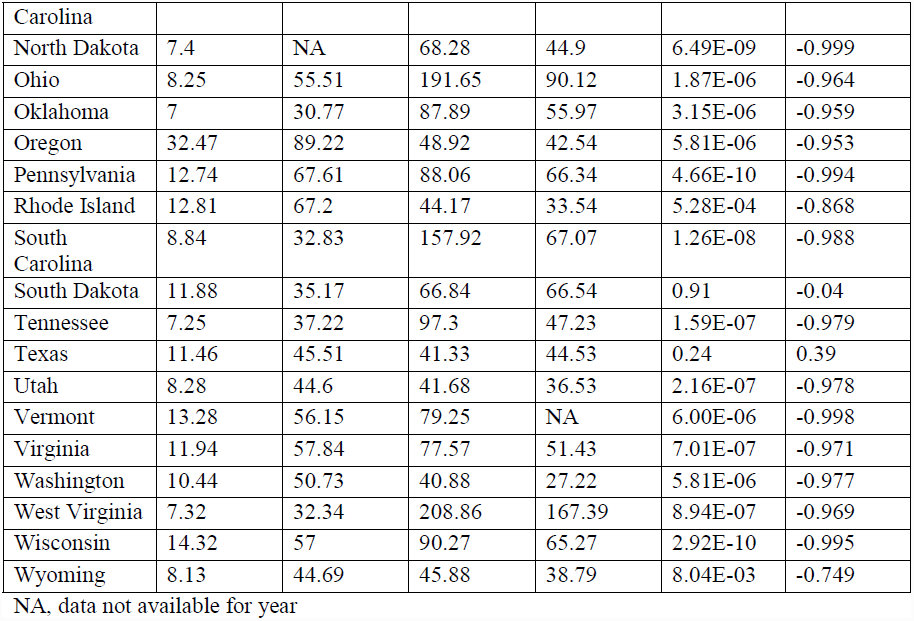
Pearson’s correlation coefficients between the prevalence of autism and that of ID for each of the 50 US states from years 2000-2010 for individuals ages 3-21 within special education.

## DISCUSSION

We analyzed one of the largest cohorts of longitudinal special education population data, through which we observed a 331% increase in the 11-year autism prevalence. This trend disappeared when the combined prevalence of autism and its frequently comorbid features were considered (Figure S2). The phenomenon of diagnostic recategorization has been noted previously^3 4^, however, the magnitude of effect from comorbid features have not been documented. Our study shows a 22-fold drop in prevalence increase when considering the prevalence of a broader neurodevelopmental disorder category including both autism and ID. Further, for children between ages 3 and 18 years, an average of 64.2% of the increase in prevalence of autism can be accounted for by a concomitant decrease in the prevalence of ID. The proportion of ID cases potentially undergoing diagnostic recategorization to autism was higher among older children (75%) than younger children (48%). These results suggest that comorbid features can confound true prevalence estimates of the autism disorder. We also found that the disability categories within the special education data showed distinct age-specific prevalence rates. For example, prevalence estimates peaked between ages 7 and 9 years for autism and between ages 11 and 18 years for ID (Figure S6). These prevalence peaks suggest distinct developmental trajectories and specific diagnostic windows for certain comorbid phenotypes. Interestingly, one recent study found that older children with autism were more likely to retain their diagnosis than those diagnosed at a younger age suggesting the complexities associated with using one set of identifiable features as diagnostic criteria^27^. It is likely that older children are more severely affected manifesting intellectual disability and other comorbid phenotypes at a later age.

In this study, we primarily focus on the comorbidity of autism and ID; however, we note that other disorders can also potentially contribute to a diagnostic recategorization to autism. For example, a significant negative correlation was observed between the prevalence of autism and that of specific learning disability (Figure S6). While we also found a significant negative correlation between the prevalence of autism and that of ID for all the US states as a whole, when assessed individually, not all states showed the same strength of correlation. For example, North Dakota, Vermont, and Georgia showed the strongest correlation coefficients (-0.999, -0.998, -0.997, respectively). However, states such as Arizona, New Jersey, and Wyoming showed less strong correlation coefficients (-0.625, -0.621, -0.749, respectively) and certain states, such as California, New Mexico, and Texas, showed no correlation at all. While the differential rates of autism prevalence reflect differences in ascertainment in special education schools across the US states, documented evidence of inconsistency in ascertainment even among groups following set diagnostic criteria suggests extensive heterogeneity of the disorder^28^. In fact, a recent study found the prevalence of autism and ID to be associated with state-related regulatory factors, and even found strong correlations between smaller, county-related factors and autism and ID prevalence^5^. Another report in 2001 found the prevalence of autism within The Brick Township, New Jersey, to be significantly higher than that of the US^23^. Further, Davidovitch and colleagues found a lower prevalence of autism among an Israeli population compared to the US^29^. A recent study in the United Kingdom using the UK General Practice Research Database showed a strikingly similar incidence of autism over a period of 10 years suggesting no apparent increase in prevalence rates^30^. These examples indicate variability in ascertainment of children with neurodevelopmental disorders across different regions, and highlight an importance for large-scale studies of autism prevalence to take these health policy variations into account.

Several clinical studies have documented varying percentages of comorbid features suggesting that comorbidity in autism is the norm rather than the exception. Changes in nosology as suggested by revisions to the DSM have certainly contributed to the deviations from the original description of autism. This is reflected by the fact that only 81.2% of children previously diagnosed with autism by DSM-IV met the criteria according to DSM-V^31^. However, studies estimating prevalence of autism seem to focus on one dimension of clinical features, often ignoring other comorbid features. For example, 40.2% of individuals identified with autism by the ADDM network were actually enrolled under eight different special education categories other than autism (Figure S7). These rates also varied among states within the ADDM network, further suggesting a major impact by state-specific policies. While it would be important to understand how these comorbidity rates change over time, one limitation of the IDEA dataset was that individuals were only placed into a single diagnostic category.

The relatively high rate of comorbidity within autism may be due to a wide array of common genes implicated in many neurodevelopmental disorders^32^. Further, core components in autism diagnosis show a documented overlap in clinical features such as language impairments^33^. Interestingly, when individuals with classically defined genetic syndromes were evaluated for autism using standardized instruments, higher frequency of autistic features were observed. In fact, some of these were never thought to be an autism disorder. For example, the frequency of autism in Smith-Magenis syndrome, a disorder characterized by severe intellectual disability/multiple congenital anomalies, was reported to be as high as 90%^34^ (Table 2). While these studies suggest that autistic features are pervasive in neurodevelopmental disorders, it is possible that many autism diagnosis instruments lose specificity when applied to severe intellectual disability disorders. These factors may create a confounding effect on autism diagnosis.

**Table 2:** Frequency of autism features in classically defined genetic syndromes.

| **Disorder** | **Frequency** | **Reference** | **Autism diagnosis instrument** |
| --- | --- | --- | --- |
| 22q11.2 deletion syndrome | 40% | Niklasson, 2009<sup>37</sup> | DSM-IV |
| Angelman Syndrome | 42% | Peters, 2004<sup>38</sup> | ADOS/DSM-IV |
| Beckwith-Wiedemann Syndrome | 7% | Kent, 2008<sup>39</sup> | Previous diagnosis |
| Charge Syndrome | 28% | Hartshorne, 2005<sup>40</sup> | ABC |
| Chromosome 2q Terminal Deletion | 24% | Casas, 2004<sup>41</sup> | Previous diagnosis |
| Cohen Syndrome | 79% | Howlin, 2005<sup>42</sup> | ADOS |
| Cornelia De Lange Syndrome | 83% | Srivastava, 2014<sup>43</sup> | CARS |
| Cowden Syndrome | 53% | Varga, 2009<sup>44</sup> | DSM-IV |
| Down Syndrome | 19% | Moss, 2013<sup>45</sup> | SCQ |
| Fragile-X Syndrome | 63% | Garcia-Nonell, 2008<sup>46</sup> | ADOS-G/DSM-IV |
| Inverted 8p Deletion Syndrome | 75% | Fisch, 2011<sup>47</sup> | CARS |
| Jacobsen Syndrome | 47% | Akshoomoff, 2014<sup>48</sup> | ADOS |
| Klinefelter Syndrome | 27% | Bruining, 2008<sup>49</sup> | ADI-R |
| Lujan–Fryns Syndrome | 63% | Lerma-Carrillo, 2006<sup>50</sup> | Unspecified |
| Moebius Syndrome | 40% | Johansson, 2007<sup>51</sup> | DSM-3R/ICD-10 |
| Neurofibromatosis Type 1 | 4% | Williams and Hersh, 1998<sup>52</sup> | DSM-IV |
| Phelan-McDermid Syndrome | 94% | Phelan, 2001<sup>53</sup> | CARS |
| Potocki–Lupski syndrome | 66% | Treadwell-Deering, 2010<sup>54</sup> | ADI-R and ADOS |
| Prader-Willi Syndrome | 36% | Lo, 2013<sup>55</sup> | DISCO |
| Smith Magenis Syndrome | 90% | Laje, 2010<sup>34</sup> | SRS, SCQ |
| Smith-Lemli-Opitz Syndrome | 71-86% | Sikora, 2006<sup>56</sup> | ADOS |
| Sotos Syndrome | 68% | Zafeirou 2013<sup>57</sup> | SCQ |
| Timothy Syndrome | 80% | Splawski, 2004<sup>58</sup> | Unspecified |
| Williams-Beuren Syndrome | 93% | Klein-Tasman 2009<sup>59</sup> | ADOS |
| Wolf-Hirschhorn Syndrome | 5% | Fisch, 2010<sup>60</sup> | CARS |

ADOS- Autism diagnostic observation schedule; ADI-R-Autism diagnostic interview-revised, ABC-Autism behavior checklist, CARS-Childhood autism rating scale, DISCO-The diagnostic interview for social and communication, SRS-Social responsiveness scale, SCQ-Social communication questionnaire, DSM- Diagnostic and statistical manual of mental disorders

In conclusion, we propose that nosologically distinct neurodevelopmental phenotypes are not necessarily independent entities and can appear during early or late developmental stages and coexist as comorbid features in an affected individual. We find that a significant proportion of individuals with autism also have a range of comorbid features, which may confound diagnosis, affecting the perceived prevalence of autism. This may be due to the emphasis given to the autism component of their diagnoses, as compared to emphasis on the comorbid features in the past years. The differences in the relative severity of each of these comorbid features can complicate definitive diagnosis. Evidently, because these features co-occur to a large extent, they transcend diagnostic boundaries and contribute to the variability and severity as well as confound disease ascertainment. It is therefore clear that the patterns of underlying genetic etiology neither map well onto current disease “models” nor respect the DSM categories^35 36^. Large-scale longitudinal studies with detailed phenotyping and deep molecular genetic analyses are necessary to completely understand the cause and effect of these “disease models”. It is important that future studies of autism prevalence take these factors into account.

## ACKNOWLEDGEMENTS

We thank Dr. Evan Eichler, Dr. Sarah Elsea, Dr. Paul Medvedev, Dr. Catarina Campbell, Dr. Karyn Meltz-Steinberg, and Dr. Francesca Chiaramonte, and members of the Girirajan lab for critical reading and comments on the manuscript. The authors declare that no conflict of interest exists in relation to this work.

## Contributors

Santhosh Girirajan conceived and designed the work. Andrew Polyak performed the analysis. Richard Kubina helped with data acquisition. Andrew Polyak and Santhosh Girirajan wrote the manuscript. Andrew Polyak, Kubina, and Santhosh Girirajan approved the final version of the manuscript.

## Web Resources

The URLs for data presented herein are as follows:

The Individuals with Disabilities Education Act database, http://www.ideadata.org United States Census Bureau: http://www.census.gov/popest/data/intercensal/

## References

1. American Psychiatric Association. Diagnostic and statistical manual of mental disorders (4th ed., Text Revision). Washington, DC: American Psychiatric Association, 2000.

2. Autism, Developmental Disabilities Monitoring Network Surveillance Year Principal I, Centers for Disease C, et al. Prevalence of autism spectrum disorders--Autism and Developmental Disabilities Monitoring Network, 14 sites, United States, 2008. MMWR Surveill Summ 2012;61(3):1–19.

3. King M, Bearman P. Diagnostic change and the increased prevalence of autism. Int J Epidemiol 2009;38(5):1224–34.

4. Shattuck PT. The contribution of diagnostic substitution to the growing administrative prevalence of autism in US special education. Pediatrics 2006;117(4):1028–37.

5. Rzhetsky A, Bagley SC, Wang K, et al. Environmental and state-level regulatory factors affect the incidence of autism and intellectual disability. PLoS Comput Biol 2014;10(3):e1003518.

6. Kim YS, Leventhal BL, Koh YJ, et al. Prevalence of autism spectrum disorders in a total population sample. Am J Psychiatry 2011;168(9):904–12.

7. Prior M. Is there an increase in the prevalence of autism spectrum disorders? J Paediatr Child Health 2003;39(2):81–2.

8. Lai MC, Lombardo MV, Baron-Cohen S. Autism. Lancet 2014;383(9920):896–910.

9. Levy SE, Mandell DS, Schultz RT. Autism. Lancet 2009;374(9701):1627–38.

10. Coe BP, Girirajan S, Eichler EE. A genetic model for neurodevelopmental disease. Curr Opin Neurobiol 2012;22(5):829–36.

11. Mitchell KJ. The genetics of neurodevelopmental disease. Curr Opin Neurobiol 2011;21(1):197–203.

12. Cristino AS, Williams SM, Hawi Z, et al. Neurodevelopmental and neuropsychiatric disorders represent an interconnected molecular system. Mol Psychiatry 2013.

13. Gadow KD. Schizophrenia spectrum and attention-deficit/hyperactivity disorder symptoms in autism spectrum disorder and controls. J Am Acad Child Adolesc Psychiatry 2012;51(10):1076–84.

14. King BH, Lord C. Is schizophrenia on the autism spectrum? Brain Res 2011;1380:34–41.

15. Rapoport J, Chavez A, Greenstein D, et al. Autism spectrum disorders and childhood-onset schizophrenia: clinical and biological contributions to a relation revisited. J Am Acad Child Adolesc Psychiatry 2009;48(1):10–8.

16. Sporn AL, Addington AM, Gogtay N, et al. Pervasive developmental disorder and childhood-onset schizophrenia: comorbid disorder or a phenotypic variant of a very early onset illness? Biol Psychiatry 2004;55(10):989–94.

17. Yeargin-Allsopp M, Rice C, Karapurkar T, et al. Prevalence of autism in a US metropolitan area. JAMA 2003;289(1):49–55.

18. Tuchman R, Rapin I. Epilepsy in autism. Lancet Neurol 2002;1(6):352–8.

19. Viscidi EW, Triche EW, Pescosolido MF, et al. Clinical characteristics of children with autism spectrum disorder and co-occurring epilepsy. PLoS ONE 2013;8(7):e67797.

20. Noh HJ, Ponting CP, Boulding HC, et al. Network topologies and convergent aetiologies arising from deletions and duplications observed in individuals with autism. PLoS Genet 2013;9(6):e1003523.

21. Steinberg J, Webber C. The roles of FMRP-regulated genes in autism spectrum disorder: single- and multiple-hit genetic etiologies. Am J Hum Genet 2013;93(5):825–39.

22. Fromer M, Pocklington AJ, Kavanagh DH, et al. De novo mutations in schizophrenia implicate synaptic networks. Nature 2014;506(7487):179–84.

23. Bertrand J, Mars A, Boyle C, et al. Prevalence of autism in a United States population: the Brick Township, New Jersey, investigation. Pediatrics 2001;108(5):1155–61.

24. Boyle CA, Boulet S, Schieve LA, et al. Trends in the prevalence of developmental disabilities in US children, 1997-2008. Pediatrics 2011;127(6):1034–42.

25. Maenner MJ, Durkin MS. Trends in the prevalence of autism on the basis of special education data. Pediatrics 2010;126(5):e1018–25.

26. Kogan MD, Blumberg SJ, Schieve LA, et al. Prevalence of parent-reported diagnosis of autism spectrum disorder among children in the US, 2007. Pediatrics 2009;124(5):1395–403.

27. Wiggins LD, Baio J, Schieve L, et al. Retention of autism spectrum diagnoses by community professionals: findings from the autism and developmental disabilities monitoring network, 2000 and 2006. J Dev Behav Pediatr 2012;33(5):387–95.

28. Lord C, Petkova E, Hus V, et al. A multisite study of the clinical diagnosis of different autism spectrum disorders. Arch Gen Psychiatry 2012;69(3):306–13.

29. Davidovitch M, Hemo B, Manning-Courtney P, et al. Prevalence and incidence of autism spectrum disorder in an Israeli population. J Autism Dev Disord 2013;43(4):785–93.

30. Taylor B, Jick H, Maclaughlin D. Prevalence and incidence rates of autism in the UK: time trend from 2004–2010 in children aged 8 years. BMJ open 2013;3(10):e003219.

31. Maenner MJ, Rice CE, Arneson CL, et al. Potential impact of DSM-5 criteria on autism spectrum disorder prevalence estimates. JAMA psychiatry 2014;71(3):292–300.

32. Pettersson E, Anckarsater H, Gillberg C, et al. Different neurodevelopmental symptoms have a common genetic etiology. J Child Psychol Psychiatry 2013;54(12):1356–65.

33. Taylor MJ, Charman T, Robinson EB, et al. Language and traits of autism spectrum conditions: evidence of limited phenotypic and etiological overlap. Am J Med Genet B Neuropsychiatr Genet 2014;165B(7):587–95.

34. Laje G, Morse R, Richter W, et al. Autism spectrum features in Smith-Magenis syndrome. Am J Med Genet C Semin Med Genet 2010;154C(4):456–62.

35. Lichtenstein P, Carlstrom E, Rastam M, et al. The genetics of autism spectrum disorders and related neuropsychiatric disorders in childhood. Am J Psychiatry 2010;167(11):1357–63.

36. Kendler KS. Advances in our understanding of genetic risk factors for autism spectrum disorders. Am J Psychiatry 2010;167(11):1291–3.

37. Niklasson L, Rasmussen P, Oskarsdottir S, et al. Autism, ADHD, mental retardation and behavior problems in 100 individuals with 22q11 deletion syndrome. Res Dev Disabil 2009;30(4):763–73.

38. Peters SU, Beaudet AL, Madduri N, et al. Autism in Angelman syndrome: implications for autism research. Clin Genet 2004;66(6):530–6.

39. Kent L, Bowdin S, Kirby GA, et al. Beckwith Weidemann syndrome: a behavioral phenotype-genotype study. Am J Med Genet B Neuropsychiatr Genet 2008;147B(7):1295–7.

40. Hartshorne TS, Grialou TL, Parker KR. Autistic-like behavior in CHARGE syndrome. Am J Med Genet A 2005;133A(3):257–61.

41. Casas KA, Mononen TK, Mikail CN, et al. Chromosome 2q terminal deletion: report of 6 new patients and review of phenotype-breakpoint correlations in 66 individuals. Am J Med Genet A 2004;130A(4):331–9.

42. Howlin P, Karpf J, Turk J. Behavioural characteristics and autistic features in individuals with Cohen Syndrome. Eur Child Adolesc Psychiatry 2005;14(2):57–64.

43. Srivastava S, Landy-Schmitt C, Clark B, et al. Autism traits in children and adolescents with Cornelia de Lange syndrome. Am J Med Genet A 2014;164A(6):1400–10.

44. Varga EA, Pastore M, Prior T, et al. The prevalence of PTEN mutations in a clinical pediatric cohort with autism spectrum disorders, developmental delay, and macrocephaly. Genet Med 2009;11(2):111–7.

45. Moss J, Richards C, Nelson L, et al. Prevalence of autism spectrum disorder symptomatology and related behavioural characteristics in individuals with Down syndrome. Autism 2013;17(4):390–404.

46. Garcia-Nonell C, Ratera ER, Harris S, et al. Secondary medical diagnosis in fragile X syndrome with and without autism spectrum disorder. Am J Med Genet A 2008;146A(15):1911–6.

47. Fisch GS, Davis R, Youngblom J, et al. Genotype-phenotype association studies of chromosome 8p inverted duplication deletion syndrome. Behav Genet 2011;41(3):373–80.

48. Akshoomoff N, Mattson SN, Grossfeld PD. Evidence for autism spectrum disorder in Jacobsen syndrome: identification of a candidate gene in distal 11q. Genet Med 2014.

49. Bruining H, Swaab H, Kas M, et al. Psychiatric characteristics in a self-selected sample of boys with Klinefelter syndrome. Pediatrics 2009;123(5):e865–70.

50. Lerma-Carrillo I, Molina JD, Cuevas-Duran T, et al. Psychopathology in the Lujan-Fryns syndrome: report of two patients and review. Am J Med Genet A 2006;140(24):2807–11.

51. Johansson M, Wentz E, Fernell E, et al. Autistic spectrum disorders in Mobius sequence: a comprehensive study of 25 individuals. Dev Med Child Neurol 2001;43(5):338–45.

52. Williams PG, Hersh JH. Brief report: the association of neurofibromatosis type 1 and autism. J Autism Dev Disord 1998;28(6):567–71.

53. Phelan MC, Rogers RC, Saul RA, et al. 22q13 deletion syndrome. Am J Med Genet 2001;101(2):91–9.

54. Treadwell-Deering DE, Powell MP, Potocki L. Cognitive and behavioral characterization of the Potocki-Lupski syndrome (duplication 17p11.2). J Dev Behav Pediatr 2010;31(2):137–43.

55. Lo ST, Siemensma E, Collin P, et al. Impaired theory of mind and symptoms of Autism Spectrum Disorder in children with Prader-Willi syndrome. Res Dev Disabil 2013;34(9):2764–73.

56. Sikora DM, Pettit-Kekel K, Penfield J, et al. The near universal presence of autism spectrum disorders in children with Smith-Lemli-Opitz syndrome. Am J Med Genet A 2006;140(14):1511–8.

57. Zafeiriou DI, Ververi A, Dafoulis V, et al. Autism spectrum disorders: the quest for genetic syndromes. Am J Med Genet B Neuropsychiatr Genet 2013;162B(4):327–66.

58. Splawski I, Timothy KW, Sharpe LM, et al. Ca(V)1.2 calcium channel dysfunction causes a multisystem disorder including arrhythmia and autism. Cell 2004;119(1):19–31.

59. Klein-Tasman BP, Phillips KD, Lord C, et al. Overlap with the autism spectrum in young children with Williams syndrome. J Dev Behav Pediatr 2009;30(4):289–99.

60. Fisch GS, Grossfeld P, Falk R, et al. Cognitive-behavioral features of Wolf-Hirschhorn syndrome and other subtelomeric microdeletions. Am J Med Genet C Semin Med Genet 2010;154C(4):417–26.

